# Topoisomerase 3α (TOP3A) Dependent Alternative Lengthening of Telomeres (ALT)

**DOI:** 10.1101/2024.11.18.624152

**Authors:** Prashant Khandagale, Yilun Sun, Sourav Saha, Liton Kumar Saha, Yves Pommier

**Affiliations:** Developmental Therapeutics Branch & Laboratory of Molecular Pharmacology, Center for Cancer Research, National Cancer Institute, NIH, Bethesda, MD 20892, USA; Department of Pharmacology, Physiology and Drug Development, University of Maryland, School of medicine, Baltimore, MD, 21201, USA

## Abstract

Alternative Lengthening of Telomeres (ALT) is a homologous recombination-dependent telomere elongation mechanism utilized by at least 10-15% of all cancers. Here we identified that the DNA topoisomerase, TOP3A is enriched at the telomeres of ALT cells but not at the telomeres of telomerase-positive (Tel) cancer cells. We demonstrate that TOP3A stabilizes the shelterin protein TERF2 in ALT cancer cell lines but not in Tel cells and that long non-coding telomere transcribed RNA (TERRA) enrichment at telomeres depends upon TOP3A. TOP3A also promotes the generation of single-stranded telomeric C-strand (ssTeloC) DNA, which is a recently discovered marker for ALT. Additionally, we found that inducing TOP3A-DNA-protein crosslinks in ALT cells suppresses TERRA enrichment as well as destabilizes TERF2. Taken together these observations uncover the unexplored functions of TOP3A at ALT telomeres and suggest the potential of developing an ALT-specific cancer therapeutic strategy targeting TOP3A.

## Introduction

To resolve the topological problems of the genome owing to the rigidity of duplex DNA, its extreme compaction in the nucleus and its processing by transcription, replication, DNA repair and recombinations, eukaryotic cells possess six different topoisomerases: TOP1, TOP1MT, TOP2A, TOP2B, TOP3A and TOP3B (1). Topoisomerase IIIα (TOP3A) belongs to the Type I topoisomerase family (1). It is primarily associated with DNA replication (2, 3) and only acts on single-stranded DNA substrates to remove hemicatenanes, precatenanes and hypernegative supercoils by reversibly cleaving the DNA backbone within single-stranded DNA segments, which allows the passage of another DNA segment through the break (4). We recently proposed that TOP3A removes the precatenanes formed behind replications fork during normal DNA replication (2, 3). In the nucleus, TOP3A acts in a complex with the Bloom helicase (BLM) and the two scaffolding proteins RMI1 and RMI2 (4). Within the BTR (BLM, TOP3A, RMI1/2) complex, TOP3A also plays a crucial role in the homology-directed recombination (HDR) pathways by resolving in a precise, error-free manner, the double Holliday junctions resulting from DNA branch migration, yielding non-crossover products (4, 5).

Telomeric DNA consists of repetitive sequences of six nucleotides TAAGGG at the ends of eukaryotic chromosomes (6). The termini of telomeres are protected from eliciting a DNA damage response and end-to-end chromosomal fusions by the shelterin complex, composed of TERF1, TERF2, TIN2, RAP1, TPP1 and POT1 (7). In normal somatic cells, the length of the telomeres decreases with each round of replication due to the inability of the classical bidirectional replication process to fully synthesize the lagging strands at the ends of chromosomes. The shortening of telomeres ultimately results in proliferative blockade and hence arrests cell growth (8). To ensure their proliferation, cancer cells most commonly maintain their telomeric length by reactivating the telomerase enzyme (8). Still, even in the absence of telomerase, a subset of cancer cells are able to maintain their telomeres via an homology-dependent recombination (HDR) mechanism known as Alternative lengthening of Telomere (ALT) (9), which has recently been related to the action of BLM on the C-rich telomeric DNA strand (10, 11). Osteosarcoma (45%) and glioblastoma (33%) are classical tumor types utilizing the ALT pathway as telomere maintenance mechanism (TMM) (12).

ALT cells display characteristic features including the recruitment of their telomeres in ALT-associated promyelocytic leukemia (PML) nuclear bodies (APBs) with accumulation of extrachromosomal partially single-stranded telomeric DNA (c-circles) and abundant telomere transcribed long non-coding RNA (TERRA) (13), which forms R-loops that are thought to promote telomeric break-induced replication (BIR) (14, 15).

The nuclear partner of TOP3A, BLM is an established key factor for maintaining telomere length and the clustering of telomeres in the APBs (10, 16–18). Recent studies proposed that the helicase activity of BLM unwinds the lagging strand substrates and forms 5’-flaps during telomere replication prior to BIR in ALT cells (10, 11). This raises the possibility that TOP3A acts at telomeres to resolve the intermediate structures formed by BLM during ALT-associated telomere recombinations (19).

In this study, we investigated the functional role of TOP3A in ALT cancer cells using single-cell protein immunofluorescence microscopy, TERF2 protein stability analyses, telomere fluorescence hybridization (FISH) and detection of c-circles (ssTeloC) and TERRA foci in ALT cells with genetic inactivation of TOP3A or overproduction of a toxic TOP3A forming TOP3A cleavage complexes (TOP3A-R364W) (2, 3). We also propose molecular mechanisms consistent with the activity of TOP3A as a DNA strand passage enzyme resolving double Holliday junctions (DHJ) and hypernegative supercoiling driven by D-loop extension.

## Results

### TOP3A is recruited to the telomeres of ALT cells

The protein composition of telomeres differs between ALT-positive (ALT) and telomerase-positive (Tel) cells. In both ALT and Tel cells, the shelterin complex proteins are localized at the telomeres. However, some proteins, such as TERT, ATRX, and dyskerin are specifically localized to the telomeres of Tel cells, while others, like PML and SLX4 are exclusively present at the telomeres of ALT cells (20). To test the presence of TOP3A at telomeres in ALT cells, we performed immunofluorescence microscopy experiments in U2OS cells co-expressing HALO-tagged TOP3A and GFP-tagged TERF2. As expected, high levels of TOP3A were found in mitochondria, which validates our technical approach (21). Most relevant here, we also observed that the HALO-tagged TOP3A foci in the nucleus co-localized with GFP-TERF2 (Figure 1A).

**Figure 1:**
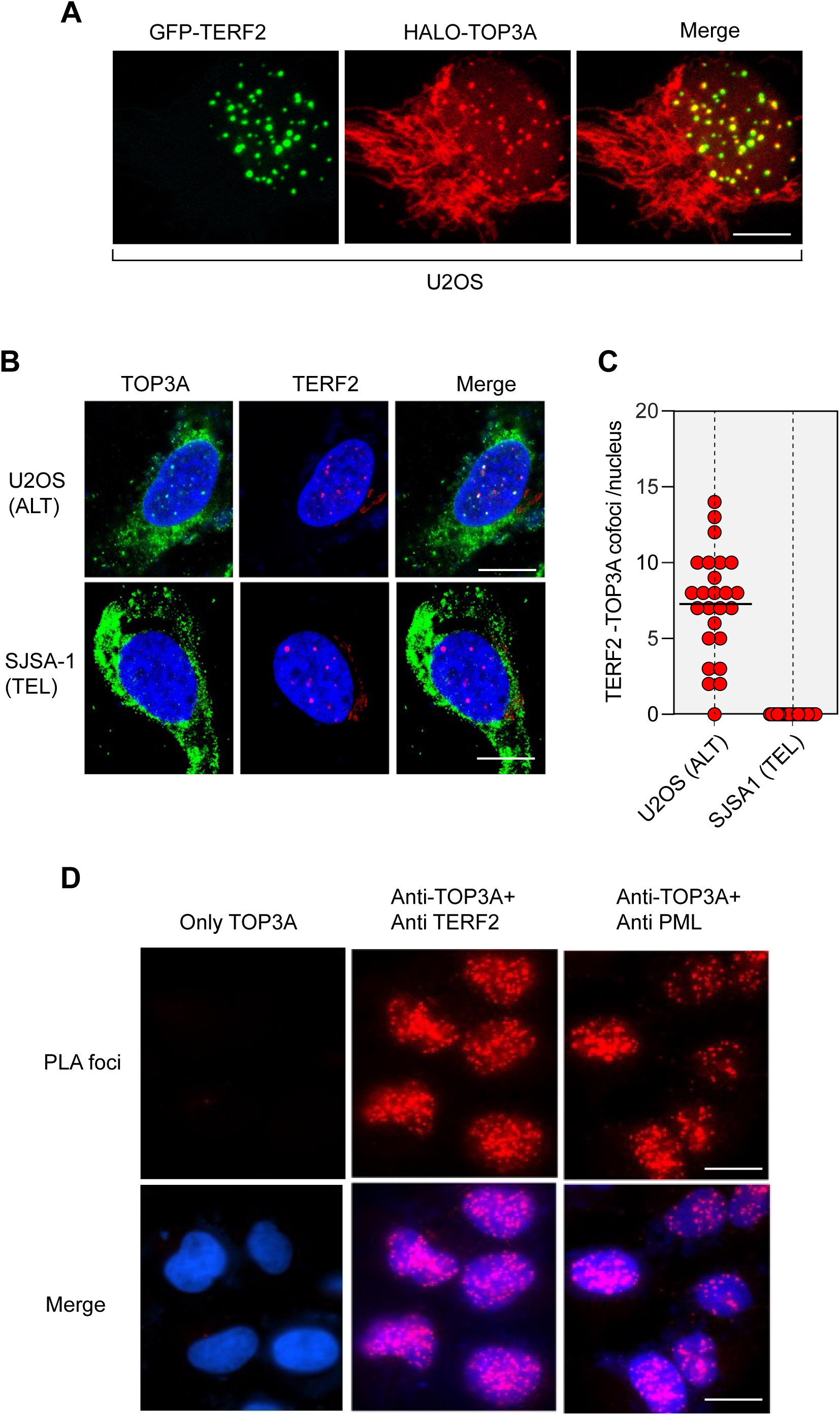
Localization of TOP3A at the telomeres of ALT cancer cells. **A.** Representative immunofluorescence images showing colocalization of HALO-TOP3A with GFP-TERF2 in U2OS. The scale bar represents 10 μm. **B.** Representative immunofluorescence images showing localization of TOP3A with TERF2 in U2OS (ALT) and SJSA1 (Tel) cells. The scale bar represents 20 μm **C.** Quantification for TOP3A-TERF2 co-foci per nucleus in U2OS and SJSA1 cells. 25 nuclei per experiment were calculated (n=2). **D.** Proximity ligation assay (PLA) showing interaction of TOP3A with TERF2 and PML in U2OS cells. The scale bar represents 10 μm.

To test the presence of endogenous TOP3A at telomeres, we performed immunofluorescence microscopy experiments with TOP3A and TERF2 antibodies both in ALT U2OS and Tel SJSA-1 cells. In U2OS (ALT) cells, TOP3A formed distinct nuclear foci in the nucleus colocalizing with TERF2 (Figure 1B and C). In SJSA-1 (Osteosarcoma Tel-positive cells (22)) (Supplementary Figure 1) (23), TOP3A was prominently detectable in mitochondria and TOP3A could not be seen in the TERF2 foci. Also, the nuclear TOP3A foci that were detectable in SJSA-1 cells tended to be small (Supplementary Figure 2). To confirm the colocalization of TOP3A and TERF2 in U2OS cells, we performed PLA assays (Figure 1D), from which conclude that TOP3A interacts closely with TERF2 in U2OS cells.

We also performed PLA assays for PML as ALT cancer cells are known to form liquid-liquid phase separation aggregates in ALT-associated PML bodies (APBs). PLA assays with TOP3A and the PML protein revealed that TOP3A also interacts with PML (Figure 1D). These experiments collectively establish the presence of TOP3A at the telomeres of ALT cells.

### APB formation is TOP3A-dependent

To determine whether lack of TOP3A affects APBs, we knocked down TOP3A and looked at APBs by dual labeling with TERF2 and PML. As expected TERF2 colocalized with PML in control cells (Figure 2A). However, TOP3A knock-down reduced the TERF2 and PML foci and their colocalization (Figure 2A-B). These results imply that the formation of APBs is TOP3A-dependent.

**Figure 2:**
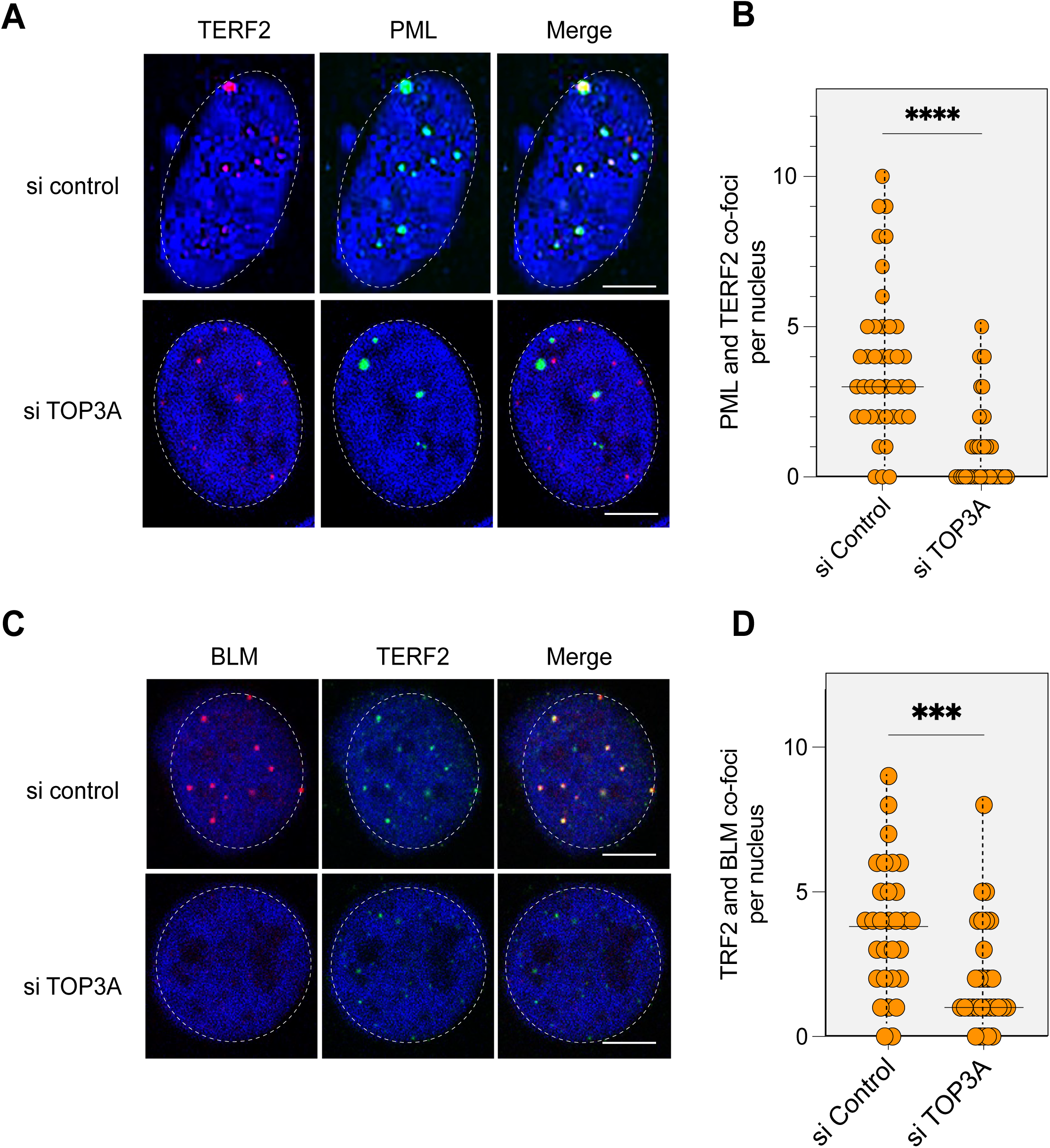
TOP3A is required for PML and BLM localization at telomeres. **A.** Representative immunofluorescence images showing localization of PML and TERF2 in U2OS cells after siTOP3A treatment. The scale bar represents 10 μm **B.** Quantification for TERF2-PML co-foci per nucleus. 40 nuclei were counted for each condition (n=3). **C.** Representative immunofluorescence images showing localization of BLM and TERF2 in U2OS cells after siTOP3A treatment. The scale bar represents 10 μm **D.** Quantification for TERF2-BLM co-foci per nucleus. 40 nuclei were counted for each condition (n=3).

As BLM helicase is an interacting partner of TOP3A and its presence at telomeres is essential for promoting the HDR-associated branch migration step (10), we tested the presence of BLM by immunofluorescence microscopy. As expected, BLM colocalized with TERF2 in control cells (Figure 2C). Notably, we also found that TOP3A knockdown reduces the recruitment of BLM to telomeres (Figure 2C-D). Together, these results demonstrate that the absence of TOP3A results in disassembly of ALT-associated PML bodies (APBs).

### TOP3A promotes the generation of ALT-associated ssTeloC DNA

ALT cells use homology-dependent recombination (HDR) mechanisms to elongate their telomeres, and during this process single-stranded segments of telomeric DNA repeats (CCCTAA) are generated due to the invasion of the broken strands into the homologous telomeres and the formation of D-loops (Figure 3A). Additionally, ALT cells contain c-circles, which contains ssTeloC DNA sequences, an established hallmark of ALT (24), (Figure 3A). As the ssTeloC signals are lost when BLM and RMI1 are inactivated in ALT cells (17), we performed native FISH assays to determine the impact of TOP3A in the ssTeloC signals. Figure 3B-C shows that knocking-down TOP3A profoundly decreases the ssTeloC signals. These results demonstrate the TOP3A is involved in the generation of the ALT-specific ssTeloC DNA signals.

**Figure 3:**
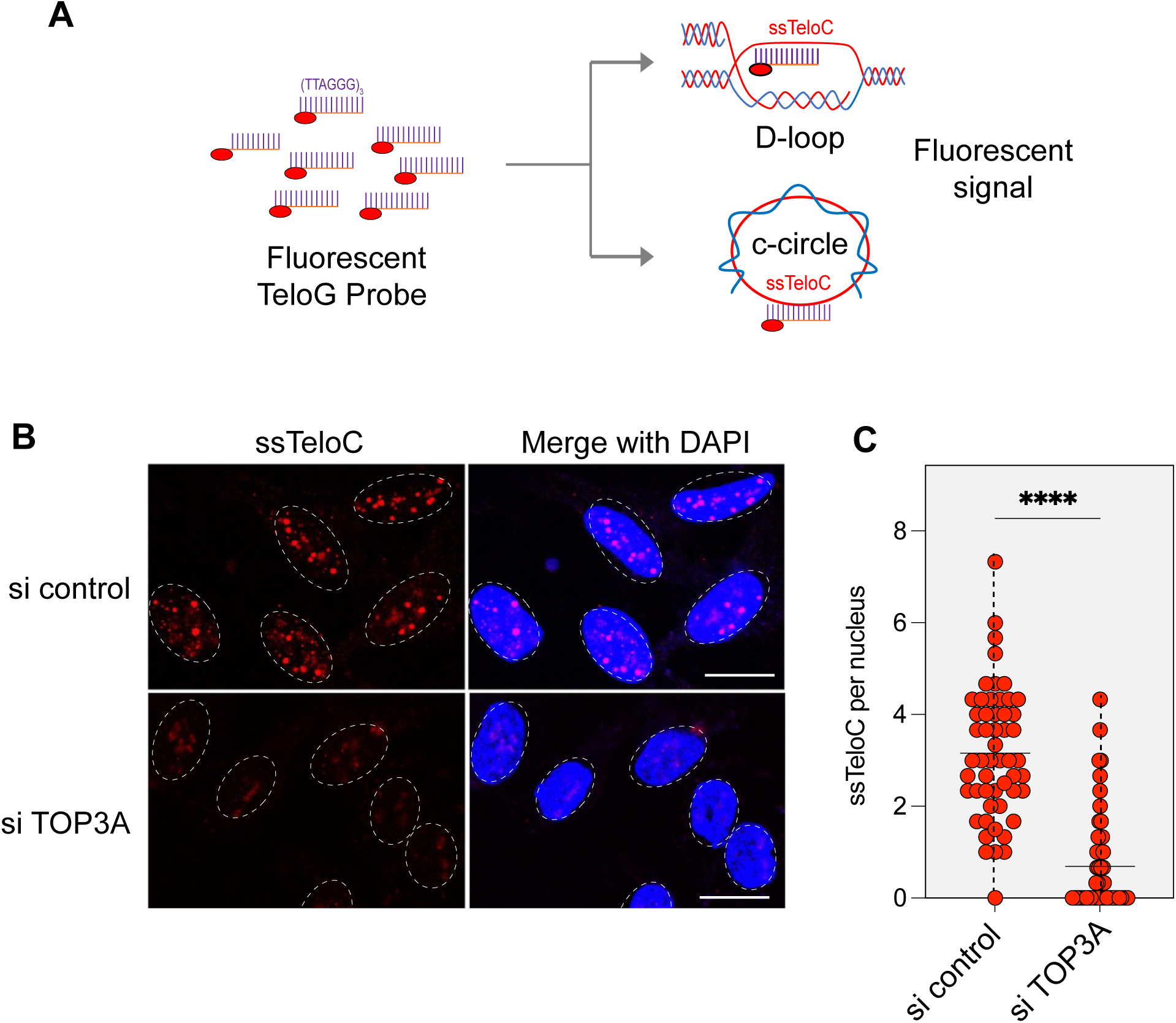
TOP3A promotes the generation of single stranded CCCTAA repeats (ss-Telo C). **A.** Schematic representation of ss-Telo C detection (telomere probe under native conditions). **B.** Representative images of ss-Telo C staining performed after native-FISH (scale bar 10 μm) **C.** Quantification for ss-Telo foci per nucleus in U2OS. 50 nuclei were counted.

### TOP3A promotes TERRA recruitment at telomeres in ALT cells

As mentioned in the Introduction, in addition to ssTeloC (see Figure 3A), non-coding telomeric RNA repeats (TERRA) represent another nucleic acid characteristics of ALT cells. TERRA is highly expressed in ALT cells, in which it can form extended RNA-DNA hybrids (R-loops; Figure 4A). To elucidate the relationship between TERRA and TOP3A, we performed RNA FISH experiments to test TERRA foci after TOP3A depletion. TERRA foci were readily detected in control cells (without RNA depletion) and these TERRA foci colocalized with TERF2 within APBs. Consistent with previous observations, these foci were sensitive to RNase A treatment (Supplementary Figure 3) (25).

**Figure 4:**
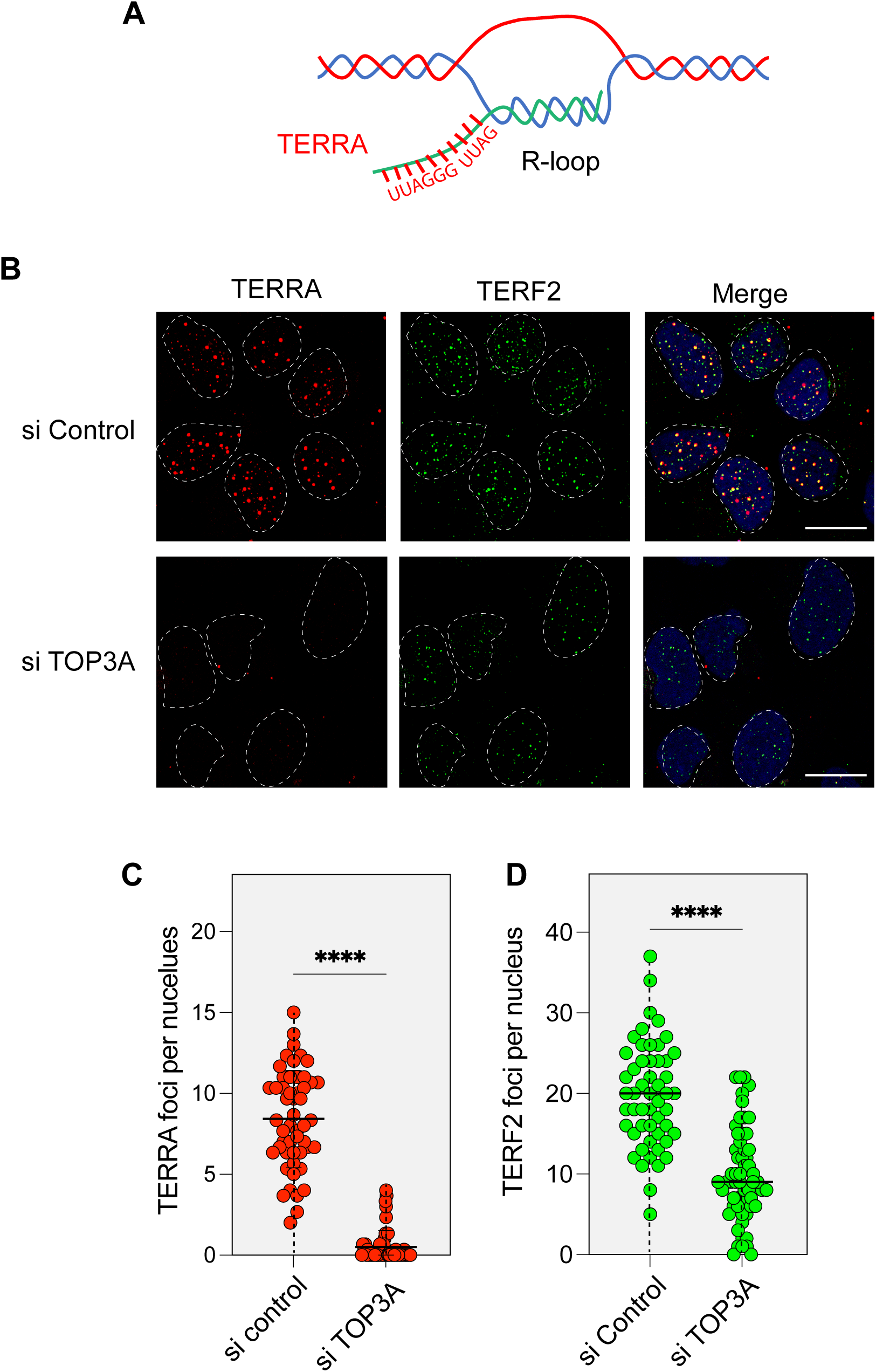
TOP3A promotes TERRA recruitment at the telomere of U2OS ALT cells. **A.** Scheme of TERRA-R-loops. **B.** Representative images of RNA-FISH for TERRA in U2OS cells after TOP3A knockdown. (Scale bar 10 μm). **C.** Quantification of TERRA foci per nucleus. 50 nuclei were counted for each condition (n = 3). **D.** Quantification for TERF2 foci per nucleus in U2OS. 40 nuclei were counted for each condition (n = 3).

Knocking-down TOP3A markedly reduced the TERRA foci (Figure 4B-C) concomitantly with a suppression of the TERF2 foci (Figure 4B-D). Of note the reduction of the TERF2 foci upon TOP3A knockdown is in agreement with the data presented in Figure 2C. From these experiments we conclude that TOP3A promotes the recruitment of TERRA in the APBs of ALT cells.

### TOP3A stabilizes the shelterin complex in ALT cells

Because genetic inactivation of TOP3A reduced TERF2 foci (see Figures 2 and 4), we examined whether TOP3A expression stabilizes the TERF2 protein. Immunoblots of cell lysates after siRNA-mediated knockdown of TOP3A were carried in both ALT and Tel cells. TERF2 protein levels were reduced after suppression of TOP3A expression in U2OS cells (Figure 5A). Two additional ALT cell lines SAOS2 and HU09 (not expressing TERT; see Supplementary Figure S4 in (23)) were tested and showed similar TERF2 depletion after knockdown of TOP3A (Supplementary Figure S3A-C). By contrast, TERF2 levels were not significantly affected by knocking down TOP3A in the TERT-positive H1080 cells (19).

**Figure 5:**
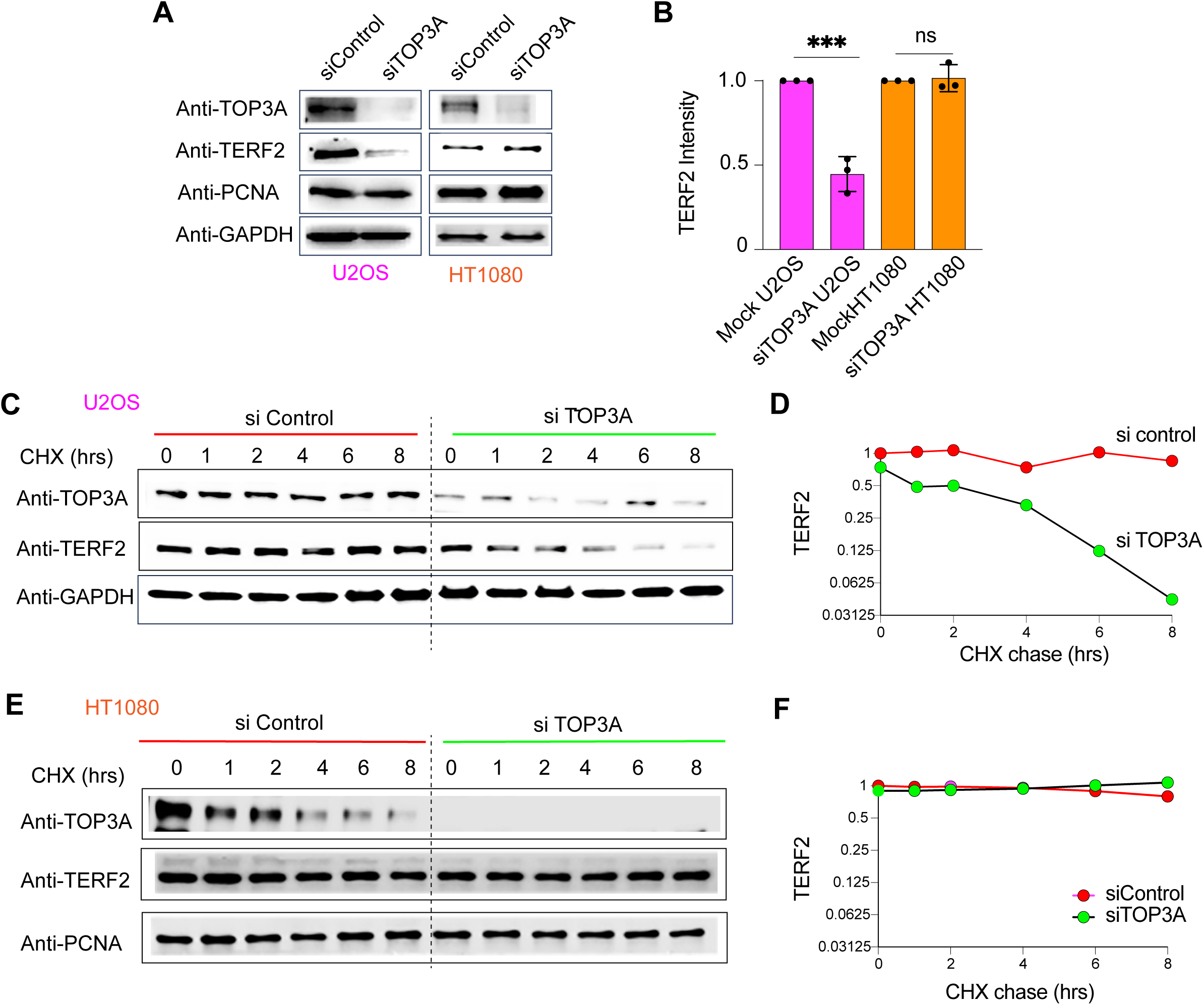
Loss of TOP3A destabilizes TERF2 in ALT cells. **A.** Representative Western blots after TOP3A knockdown in U2OS and HT1080 cells. **B.** Quantification of TERF2 intensity using image J software and normalization with GAPDH controls. The experiment was performed in triplicates and average TERF2 intensity calculated. **C.** Cycloheximide chase experiment monitoring TERF2 stability after TOP3A knockdown in U2OS cells at the indicated time intervals. **D.** Quantification of TERF2 intensity using image J software and normalization with GAPDH. **E.** Same as panel E in HT1080 cells. **F.** Quantification of TERF2 intensity using image J software and normalization with PCNA.

To test whether TOP3A acts by stabilizing TERF2 in ALT cells, we measured TERF2 protein levels in U2OS and HT1080 cells treated with cycloheximide (CHX) to suppress protein synthesis. The cycloheximide chase experiments showed time-dependent depletion of TERF2 protein in the U2OS cell treated with siTOP3A compared to cells treated with control siRNA (Figure 5C-D). In contrast, similar experiment performed in HT1080 cells showed that TERF2 levels remained unaltered after CHX treatment in both control and TOP3A depleted cells (Figure 5E-F). Together, these results demonstrate that TOP3A stabilizes TERF2 in ALT cells.

### Loss of TOP3A causes growth retardation and telomere aberrations in ALT cells

Because unprotected and destabilized telomeres should be deleterious, we compared the effect of TOP3A downregulation on the growth of U2OS (ALT) and HT1080 (Tel) cells. Figure 6A shows that knocking-down TOP3A reduces the growth of U2OS cells while having minimal effect on the HT1080 cells. Growth retardation upon TOP3A downregulation was also observed in the SAOS2 cell line (Supplemental Figure S4D). These results are consistent with a prior study showing that TOP3A preferentially sustains the growth of ALT cells (19).

**Figure 6:**
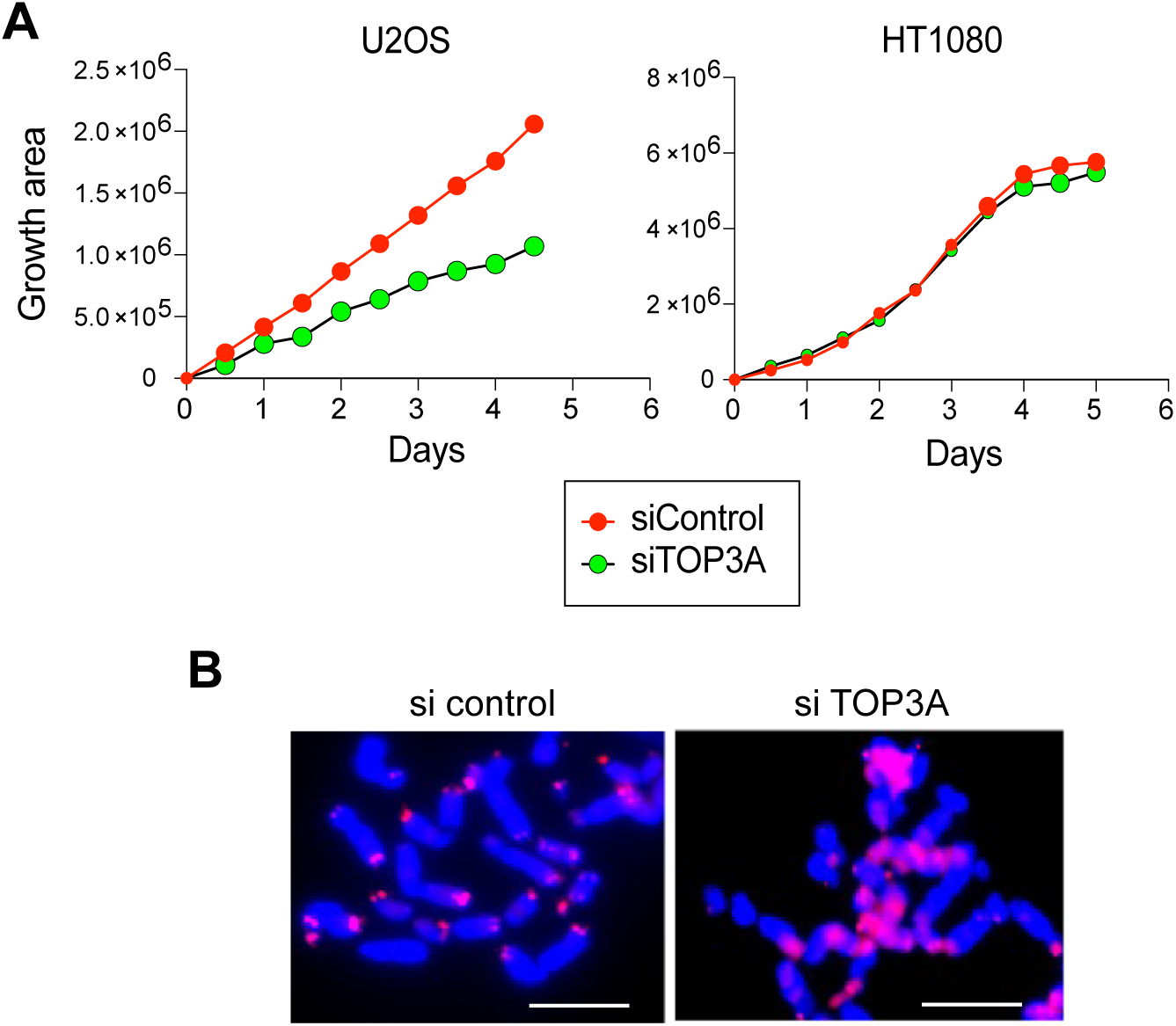
Loss of TOP3A causes telomere fragility and affects the proliferation of ALT cells. **A.** Growth of U2OS and HT1080 cells was monitored after knockdown of TOP3A using Bio-spa. The experiment was done in duplicate. **B.** TOP3A knockdown cells examined by metaphase-FISH. Telomeres were stained using CY-5 labeled telomere probe (red) and DAPI was used to stain chromosomes (blue). Scale bar 20 μm.

Because we observed loss of TERF2, a major component of the shelterin complex, in the absence of TOP3A (see above), and because absence of TERF2 results in end-to-end telomere fusions (26), we analyzed metaphase spreads coupled with telomere FISH in TOP3A-depleted U2OS cells. Figure 6B shows that upon TOP3A depletion, chromosomes displayed fragile, smeared and ultrabright telomeres. These experiments show that TOP3A is required for telomere protection in ALT cancer cells and for cellular proliferation.

### Poisoning TOP3A suppresses TERRA and TERF2 foci in ALT cells

To determine whether poisoning of TOP3A could also affect ALT telomeres, we took advantage of our recent observation that mutating a conserved arginine residue of TOP3A to tryptophane (R364W) in close proximity of TOP3A’s catalytic tyrosine (Figure 7A) prevents the enzyme from resealing the breaks it generates during its enzymatic catalytic cycle, thereby producing break-associated TOP3A DNA-protein crosslinks (DPCs) (2, 3). Overexpression of the TOP3A R364W mutant in U2OS cells showed reduced TERRA foci compared to WT TOP3A (Figure 7B-C), suggesting that even in cells expressing endogenous TOP3A, poisoning TOP3A can negatively affects the telomeres of ALT cells.

**Figure 7:**
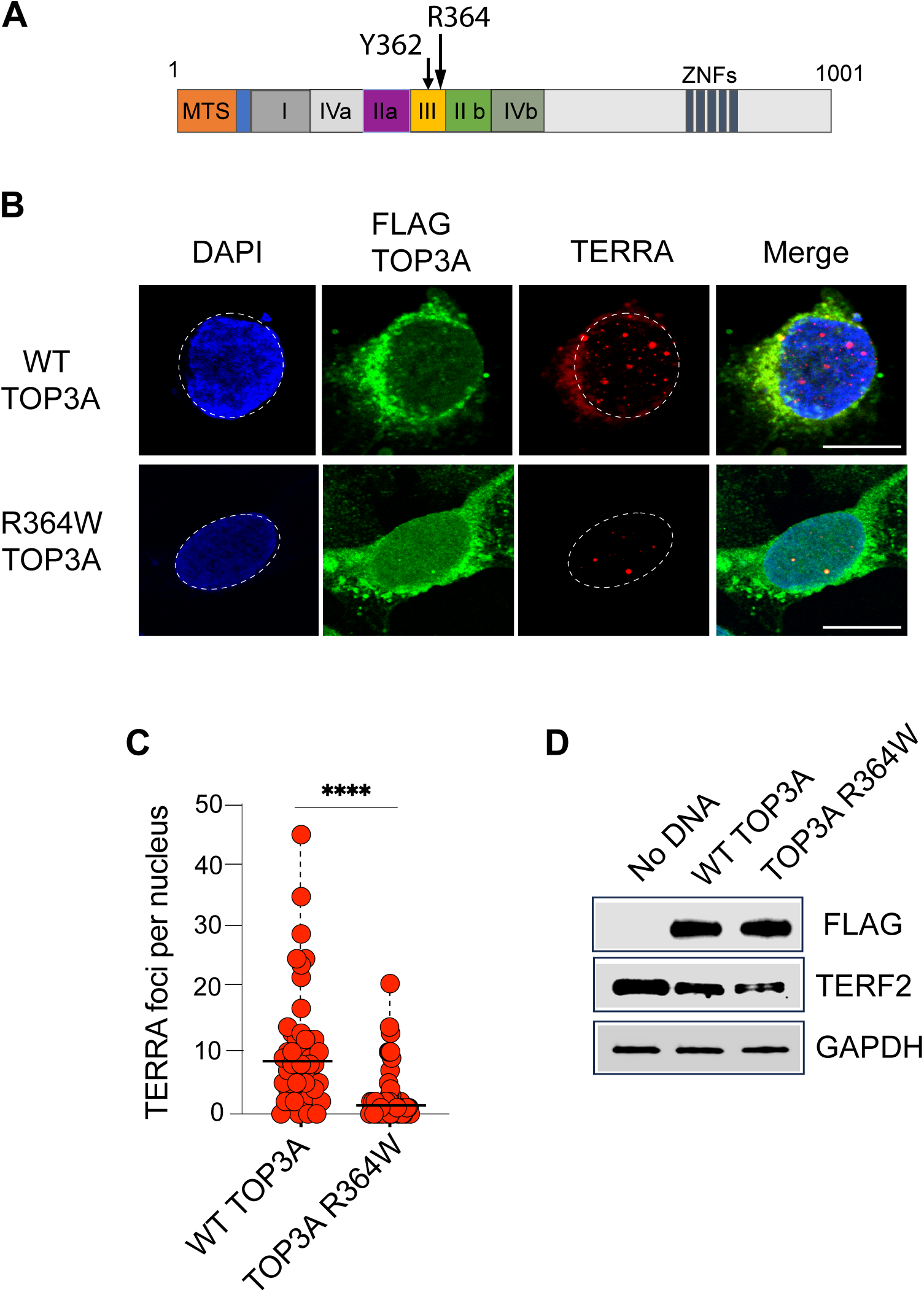
TOP3A DNA crosslinking impairs TERRA recruitment and destabilizes TERF2. **A.** Domain organization of human TOP3A. The arrow indicates the catalytic tyrosine and the adjacent arginine residue. **B.** Representative images of RNA-FISH for TERRA in U2OS cells after overexpression of WT TOP3A and TOP3A R364W. Scale bar 20 μm **C.** Quantification for TERRA foci per nucleus in U2OS cells after overexpression of WT TOP3A and TOP3A R364W. 50 nucleus per experiment were calculated (n=3). **D.** Representative TERF2 immunoblot after overexpression of WT TOP3A and TOP3A R364W.

To establish whether TOP3A dysfunction also affects TERF2, we perform immunoblotting experiments with lysates from U2OS cells transfected with WT or R364W TOP3A. While overexpression of TOP3A WT caused a mild decrease in TERF2 levels, further reduction was observed by overexpressing the TOP3A-R364W self-poisoning mutant (Figure 7D). These results suggest that poisoning TOP3A and generating TOP3A-DPCs can destabilize TERF2 and reduce TERRA foci at the telomeres of ALT cells.

## Discussion

A number of studies have unambiguously implicated the BLM helicase (10, 17, 27), to which TOP3A is coupled in the BTR complex (encompassing BLM, TOP3A, RMI1/2) (4) for telomere maintenance in ALT cells, but evidence and mechanistic insight of the implication of TOP3A have remained limited (19, 28). Our study demonstrates the direct implication of TOP3A as a critical TMM (telomere maintenance mechanism) of ALT cells, and Figure 8 summarizes our key findings and mechanistic details suggesting how TOP3A could act as telomeric topoisomerase in ALTcells.

**Figure 8:**
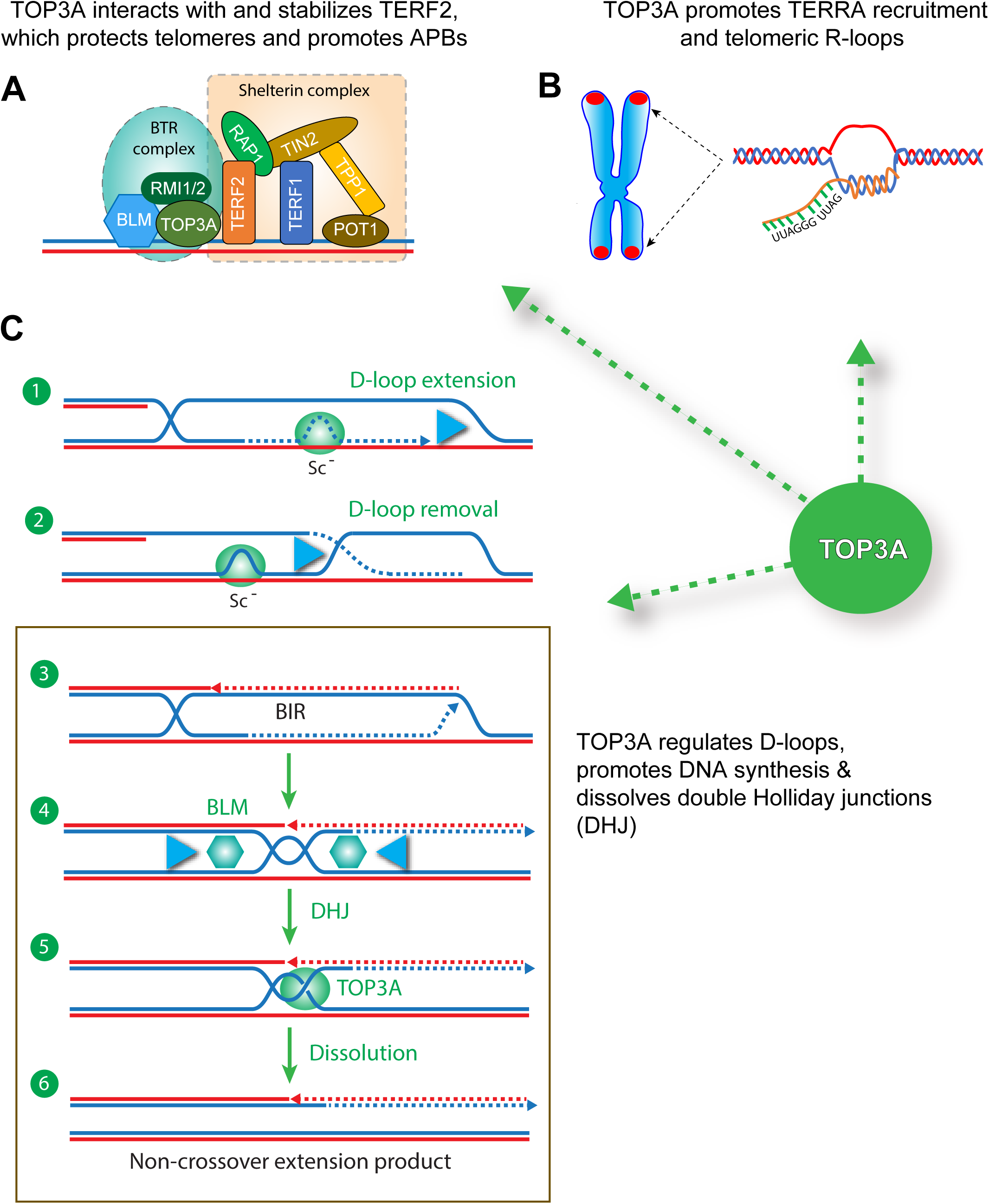
Proposed roles of TOP3A in ALT. **A.** TOP3A interacts with the components of the TRR complex and protects TERF2 and the shelterin complex from destabilization in ALT cells. **B.** TOP3A promotes the formation of TERRA R-loops and c-circles at the telomeres of ALT cells. **C.** TOP3A resolves topological DNA structures associated such as D-loops during ALT synthesis (hypernegative supercoils [Sc-] [steps 3-4]. TOP3A can dissolve double Holliday junctions (DHJ) to allow the release of extended telomeres without crossover [steps 5-6]) (see Discussion for details).

Our conclusions that TOP3A forms telomeric foci in ALT cells (Figs. 1, S2 and 7) and that ALT cells are dependent on TOP3A for maintaining their telomeres and their proliferation (Fig. 6) are consistent with a prior study (19). In addition, we provide evidence for the critical role of TOP3A for: 1/ enabling the formation of APBs as measured by single-cell analyses and immunofluorescence staining of PML and TERF2 (Fig. 2); 2/ enabling the formation of c-circles (Fig. 3) and the recruitment of TERRA at telomeres (Fig. 4), which has been proposed to initiate break-induced replication (BIR) (15, 29); and 3/ preventing the degradation of TERF2 specifically in ALT cells (Fig. 4). We also show that TOP3A does not co-localize with TERF2 in the TEL cell line SJSA-1 (Fig. 1) and that poisoning TOP3A with a genetically altered TOP3A mutation (R364W) (2, 3) is detrimental to ALT cells (Fig. 7).

Our study adds ALT-TMM (ALT-telomere maintenance mechanism) to the other cellular functions of TOP3A (4). Indeed, TOP3A is also critical for mitochondrial DNA replication (30–32) and for canonical nuclear replication before the G2/M transition to decatenate double-Holliday junctions (4, 33) and to compact DNA (34), and during S-phase presumably to avoid the accumulation precatenanes behind replication forks (2, 3, 35). Yet, we find that unless cells are ALT, TOP3A inactivation can be tolerated in cancer cells (Figs. 6 and S4), which agrees with two independent reports (19, 36).

Our finding that TOP3A deficiency leads to reduced TERF2 protein levels in ALT cells (Figs. 5 and S4) is consistent with the prior observations of Riou, Amor-Gueret and coworkers (19). We believe it is related to fact that TOP3A is a tight component of the BTR complex (4), which is itself also a prominent component of larger protein-nucleic complexes including the shelterin complex (Figure 8A) comprised in the APBs (ALT-associated PML bodies) (9, 10, 17, 19, 27). Our experiments show that TERF2 is degraded in the absence of TOP3A in ALT but not in Tel cells (Fig. 5), which leads us to propose that insufficient levels of TOP3A protein destabilize the shelterin complex and the architecture of the ABPs in ALT cells (Fig. 2) (Figure 8A). The requirement of TOP3A for the recruitment of TERRA at telomeres (see Fig. 4) in ALT cells could be related to the direct binding of TERRA to proteins of the shelterin complex and to the binding of TERRA to the C-rich strands of telomeric DNA to form R-loops (see Fig. S3) (Figure 8B).

Because TOP3A is a Type IA topoisomerase that can only act at single-stranded segments of DNA, we propose that TOP3A catalytic functions support ALT-associated BIR (break-induced replication) and the dissolution of replicated telomeric junctions during ALT. Figure 8 proposes some key steps in which TOP3A catalytic mechanisms may be required. First, TOP3A-dependent recruitment of TERRA at telomere could favor TERRA R-loop formation (Figure 8C1) to promote strand invasion and subsequent D-loop formation, as proposed by Zou and coworkers (29). Second, the ability of TOP3A to remove hypernegative supercoils (37) is a plausible site of action behind DNA tracking helicase/polymerases that either extend or remove D-loops (Figure 8C1-2) (38). Third, the enzymatic topoisomerase activity of TOP3A within the BTR complex could release ALT replication intermediates into non-crossover products by dissolving double Holliday junctions (DHJ) resulting from the convergence of the D-loop ends (16, 17, 27, 39) as it is known for TOP3A-mediated dissolution of recombination intermediates (4) (Figure 8C3-6).

Although ALT has been recognized as a rationale target for ALT cancers such as osteosarcomas, rhabdomyosarcoma (23), neuroblastomas or glioblastomas, and in spite of the increasing number of patients identified with ALT cancers (40) based on the emergence of novel genomic assays such as the DNA C-circle assay (41), no such therapies have been successfully developed (9, 42). Our findings that TOP3A inactivation selectively destabilizes TERF2 and suppresses the growth of ALT cancer cells suggest that targeting TOP3A could be a potential approach to treat ALT cancers. Because the other topoisomerases, human TOP1, TOP2A and TOP2B and the bacterial topoisomerases Gyrase and Topo IV are successfully targeted by widely used anticancer and antibacterial drugs that poison the enzyme cleavage complexes (43–45), it is plausible that drugs trapping TOP3A cleavage complexes would prove effective as anticancer agents. Our finding that the self-poisoning R364W-TOP3A was able to downregulate TERF2, APBs and TERRA foci (Fig. 7) sets of proof-of-principle for this approach.

## Materials and methods

### Reagents used

A complete list of reagents used is in Supplementary Table S1

### Cell culture, reagents and transfections

Cell lines were purchased from ATCC. U2OS, SAOS2, HT1080 cells were cultured and maintained in DMEM medium containing 10% FBS. HU09 and SJSA-1 cells were maintained in RPMI1640 medium. siRNA for TOP3A, BLM and RMI1 were purchased from horizon discovery. The siRNA transfection was performed using RNAi max transfection kit from Invitrogen according to manufactures manual. Plasmid transfections were performed using lipofectamine agent.

### Immunofluorescences for TERF2 with PML or BLM

The immunofluorescences protocol was performed as described (17). The cells were grown on coverslip were fixed with 4% paraformaldehyde for 10 minutes followed by incubation for 30 minutes in BSA blocking buffer. Primary antibodies for TERF2 (1:1000 dilution, NB110-57130, Novus Biologicals, rabbit and 1:250 dilution,4A794, Millipore sigma, mouse), PML (1:250 dilution, sc-966, Santa Cruz Biotechnology, mouse) and BLM (1:250 dilution, A-300-110, Bethyl laboratories, Rabbit) in the blocking buffer were incubated for 1 hour at room temperature. After washing the coverslips with PBS, incubation with secondary antibodies (Alexa 488, A11034 and Alexa 568, A11031 from thermos-fisher scientific) in blocking buffer for 30 minutes. Cells were washed for 3 times in the PBS and were mounted on cover glass using vectashield antifade mounting medium with DAPI. Images were captured using Zeiss LSM 780 confocal microscope.

### IF-RNA FISH

Cells grown on coverslips were washed for 30 seconds with cyto buffer 100 mM NaCl, 300 mM sucrose, 3 mM MgCl2, 10 mM pipes pH6.8) for 30 seconds, washed in cyto buffer with 0.5% Triton X-100 for 30 seconds, washed in cryobuffer for 30 second, then fixed for 10 min in 4% paraformaldehyde in PBS. Followed the cells were washed with 0.5% PBST buffer for 10 minutes. The immunofluorescences were performed with Anti-TERF2 antibody as described above. The cells were dehydrated in 70%, 90% and 100% ethanol, air-dried and hybridized overnight at 37°C with a telomere specific PNA-Cy-3 labelled Tel C (CCCTAA)_n_ probe in hybridization buffer (2× sodium saline citrate (SSC)/ 50% formamide). For control, cells were treated with RNase A (50ug/ml) for 60 minutes prior to hybridization. Coverslips are washed 3 times with hybridization buffer for 5 minutes, 3 times with 2X SSC, 1 time with 1X SSC at room temperature. The cells were mounted using vectashield antifade mounting medium with DAPI. Signals were visualized in confocal microscope zeiss 780 LSM.

### ssTelo-C staining

siRNA-treated cells were grown on coverslips and fixed with 4% paraformaldehyde for 5 minutes. The protocol was performed as described (17). In brief, cells were treated with RNase A (500ug/ml) in blocking solution (1 mg/mL BSA, 3% goat serum, 0.1% Triton X-100, 1 mM EDTA in PBS) for 1 hour at 37°C. Coverslips were dehydrated in an ethanol series (70%, 90%, and 100%) and hybridized at room temperature with a Cy-3 labelled Tel G (TTAGGG)_n_ probe (PNA Bio) in hybridization buffer (70% formamide, 1 mg/mL blocking reagent, 10 mM Tris-HCl at pH 7.2). Following the hybridization, coverslips were washed twice with buffer containing 70% formamide and 10 mM Tris-HCL in PBS and followed by 3 washes of PBS. The cells were mounted using vectashield antifade mounting medium with DAPI. Images were captured using Zeiss LSM 780 confocal microscope.

### Immunofluorescence imaging for TOP3A with TERF2

Cells grown on coverslips were serially washed 30 seconds with cyto buffer (100 mM NaCl, 300 mM sucrose, 3 mM MgCl2, 10 mM pipes pH 6.8) for 30 seconds, washed in cyto buffer with 0.5% Triton X-100 for 30 seconds, washed in cytobuffer for 60 seconds, and then fixed for 10 min in 4% paraformaldehyde in PBS, following the cells were incubate for 10 minutes in PBST. The immunofluorescence assays were performed using Anti-TERF2 (1:250) and Anti-TOP3A (1:1000) (a kind gift from Dr Jean-Francois Riou).

### Western blotting

Cells were grown in humified CO2 incubator at 37 ^O^C up to 70% confluency. After 48-72 hours of siRNA transfection the cells were harvested and washed with PBS, incubated with NTEN buffer and kept for 1 hour at 4°C in rocking condition. After centrifugation at 10,000 rpm, the supernatant was collected, and protein concentration was determined with the Bradford method. 30 ug of protein were loaded on each well of 4-12% SDS-PAGE gels.

### PLA assays

Assays were performed as described (46). In brief, the Duolink PLA fluorescence assay (Sigma-Aldrich, catalog no. DUO92101) was performed following the manufacturer’s instruction. Briefly, U2OS cells were seeded on coverslips, fixed for 15 min at 4°C in 4% paraformaldehyde in PBS and permeabilized with 0.25% Triton X-100 in PBS for 15 minutes at 4°C. The coverslips were blocked with Duolink-blocking solution and incubated with the indicated antibodies (Anti TOP3A with Anti TERF2 or Anti PML) in the Duolink antibody diluent overnight, followed by incubation with PLUS and MINUS PLA probes, ligation, and amplification. Coverslips were then washed and mounted with using mounting medium with DAPI. Images were captured on Zeiss LSM 780 confocal microscope.

### Cloning of HALO-tag TOP3A

The TOP3A-HaloTag vector was constructed with the Gateway cloning method (46). Briefly, the following primer sequences (F.P: atcgCGTCTCGGGCTGCCACCATGATCTTTCCTGTCGCCCGCTAC and R.P: atcgCGTCTCGGGGTTCAGCCGGAAATCTCGAGCGTC) were used to construct TOP3A cDNA on the Entry cloning vector. LR reaction was performed between the Entry cloning vector and the backbone vector to recombine TOP3A into the final vector backbone.

### Imaging of TOP3A-HaloTag and TERF2-GFP

The TOP3A-HaloTag and TERF2-GFP vectors were transfected into U2OS cells, using Lipofectamine 3000 following manufacturer’s instructions. After 8 hours transfection, the transfected cells were trypsinized, and 1 × 10^5^ cells in 250 μl medium were seeded in each well of the μ-Slide 8-Well plate (Ibidi). Following another 16 hours incubation, Janelia Fluor HaloTag ligand 549 was added to the wells at a final concentration of 5 nM for 30 minute incubations. The medium was removed, and the cells were washed with 1× PBS prior to the addition of 250 μl live cell imaging solution (Invitrogen) in each well. Images were captured on an instant structured illumination microscope (iSIM)(47), processed and analyzed using ImageJ.

### Metaphase spread combine FISH

Telomeric FISH on metaphase spreads were performed as described (48). In brief, cells were treated with 0.2 μg/ml colcemid 4 hours after siRNA treatment. Cells were treated harvested by trypsinization and washed with PBS, swelled in 75 mM KCl (75 mM), followed by fixation using methanol:acetic acid (3:1) solution, and metaphases were spread on glass slides. Following hybridization with an AlexaFluor 564-TelC (TAACCC) PNA probe (PNA Bio) and DNA counterstain (DAPI) images were captured on Zeiss microscope.

### Statistical analyses

Western blot quantifications were done using Image J software. *p*-value were calculated using paired Student’s t-test (two-tailed) for independent samples. ∗∗∗∗, indicates a statistical significance computed p value <0.0001, ∗∗∗ indicates p value <0.001 and n.s. = not significant.

## Acknowledgements

We thank microscopy core facility, Center for Cancer Research (CCR), NCI, NIH for imaging. We are also thankful to Shar-yin Huang for helping with the Bio-spa instrument. We are grateful Jean-Francois Riou, Muséum d’Histoire Naturelles, Paris, France, for the TOP3A antibody used in IF experiment. This study was supported by the Center for Cancer Research, the Intramural Program of the National Cancer Institute, National Institutes of Health, Bethesda, Maryland 20892 (BC 006150).

## Author contributions

P.K. and Y.P. designed research; P.K., Y.S., S.S. and L.K.S. performed research; P.K. and Y.P. analyzed data; and P.K., Y.S., S.S., L.K.S. and Y.P. wrote the manuscript.

---

**Supplementary Figure S1:**
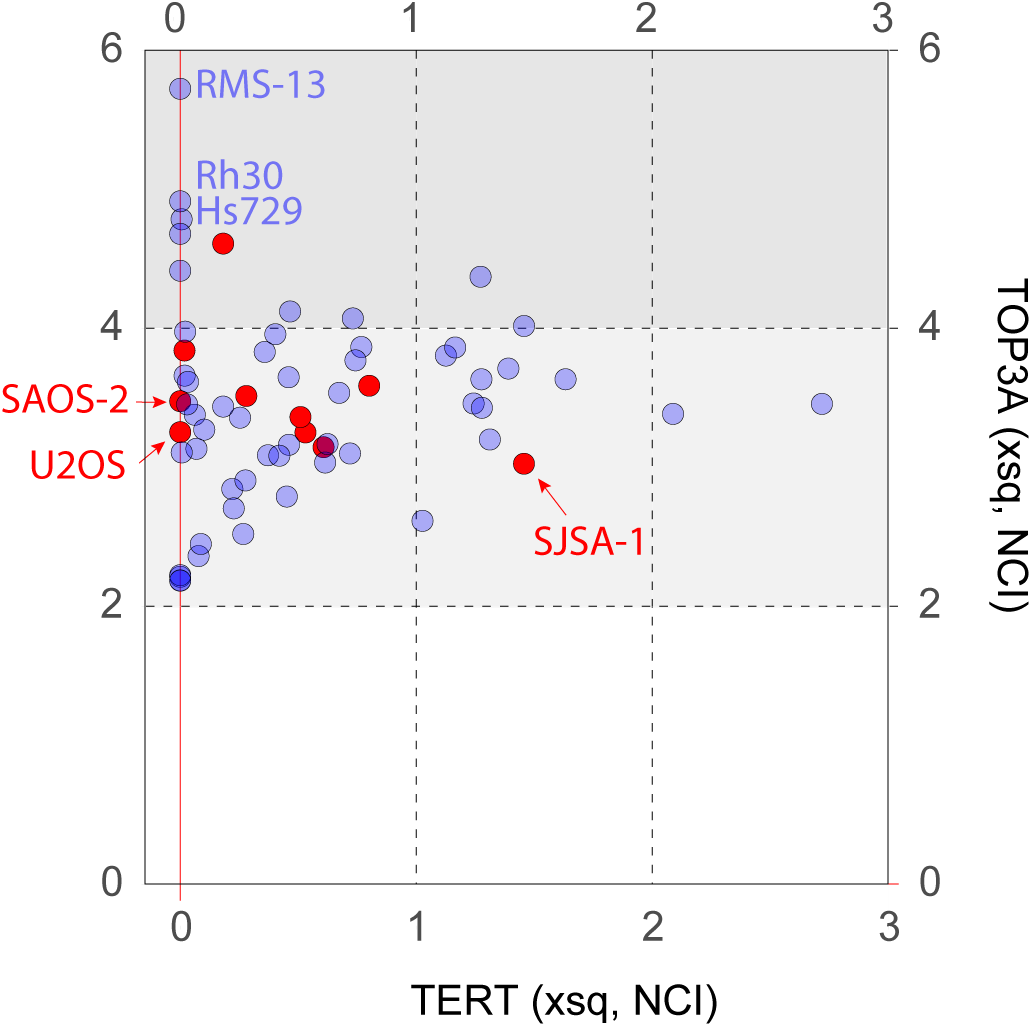
Snapshot image of Sarcoma CellMinerCDB (https://discover.nci.nih.gov/) (23) confirming lack of telomerase (TERT) expression in the U2OS and SAOS-2 cell lines while the SJSA-1 cell lines (ALT negative) express TERT. Each dot corresponds to a sarcoma cell line from the NCI database with the osteosarcoma cell lines in red. Note that all cell lines express TOP3A and that the three TERT-negative rhabdomyosarcoma cell lines (RMS-13, Rh30 and Hs729) show high TOP3A expression (above 4 in the area shaded in gray; units on the axis are Log_2_FPKM +1).

**Supplementary Figure S2:**
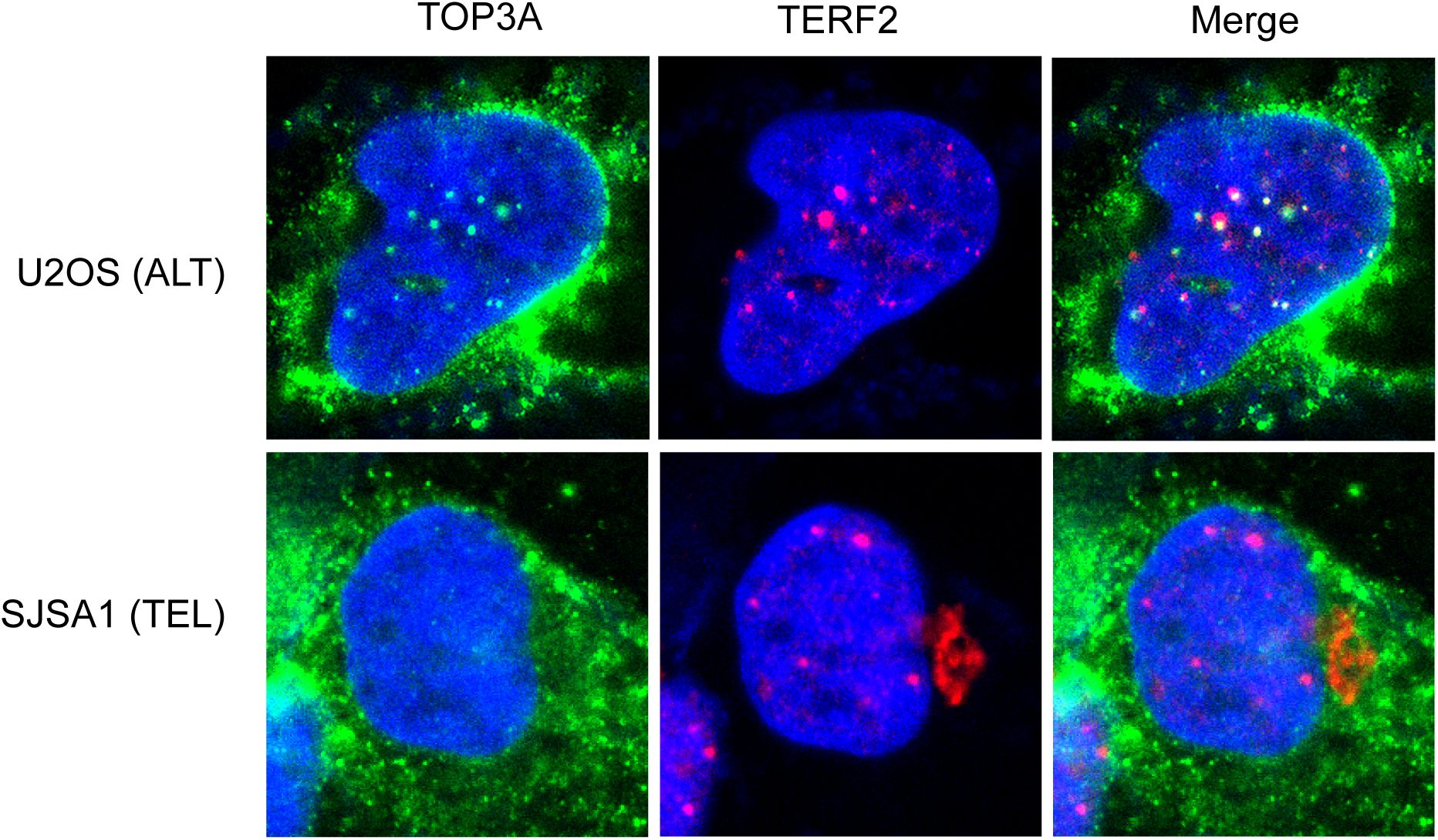
Representative immunofluorescence microscopy images showing that nuclear TOP3A foci colocalize with TERF2 foci in a U2OS (ALT) cell. By contrast, in SJSA1 cells (TEL-positive) TOP3A nuclear foci were barely visible and were not detectable at TERF2 foci. As expected, TOP3A staining is most prominent in mitochondria.

**Supplementary Figure S3:**
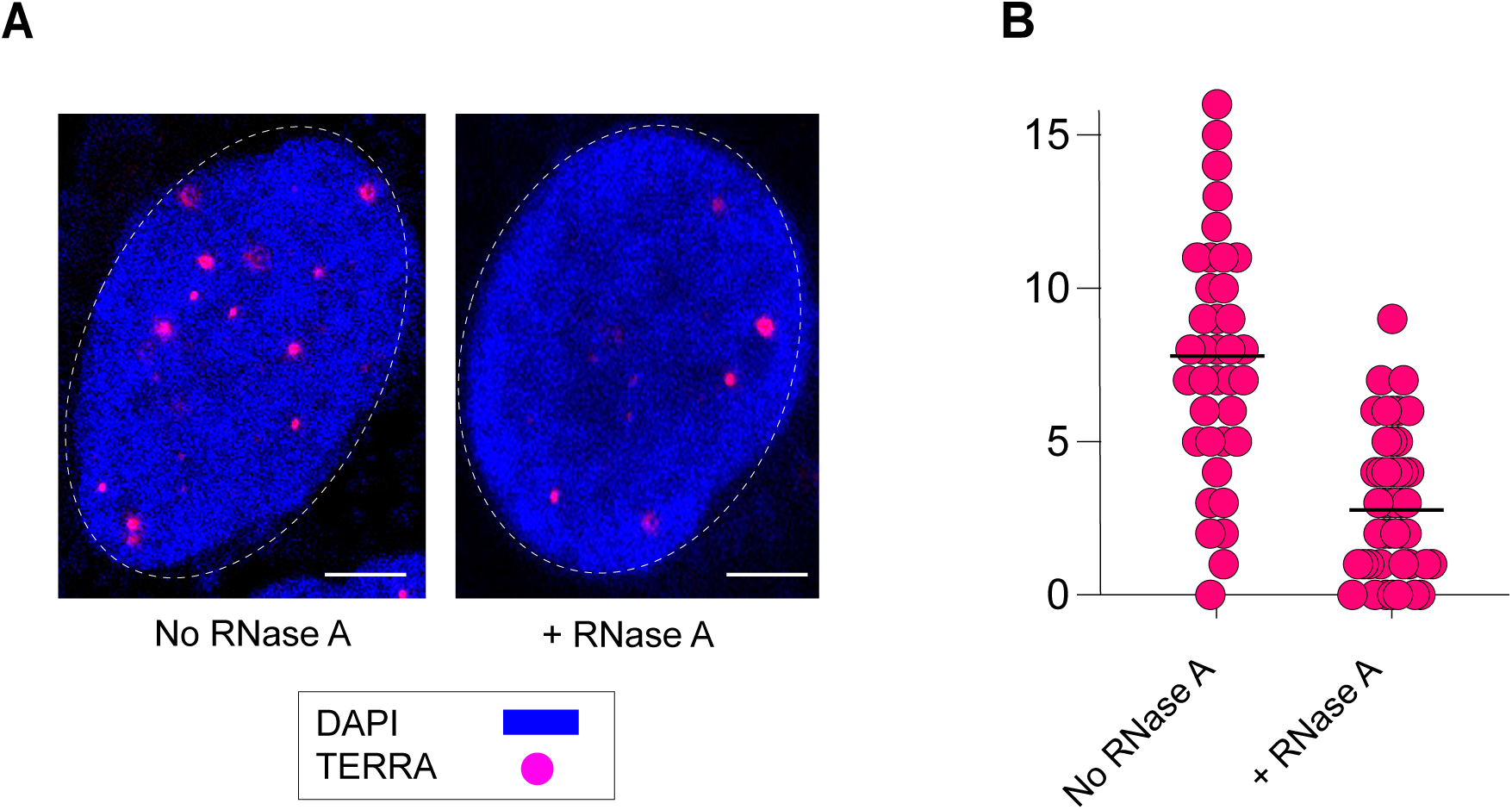
A. Representative images of ss-Telo G staining performed in U2OS cells with and without RNase. Scale bar 10 um. **B.** Quantification for ss-telo G foci per nucleus. 30 nuclei were counted.

**Supplementary Figure S4:**
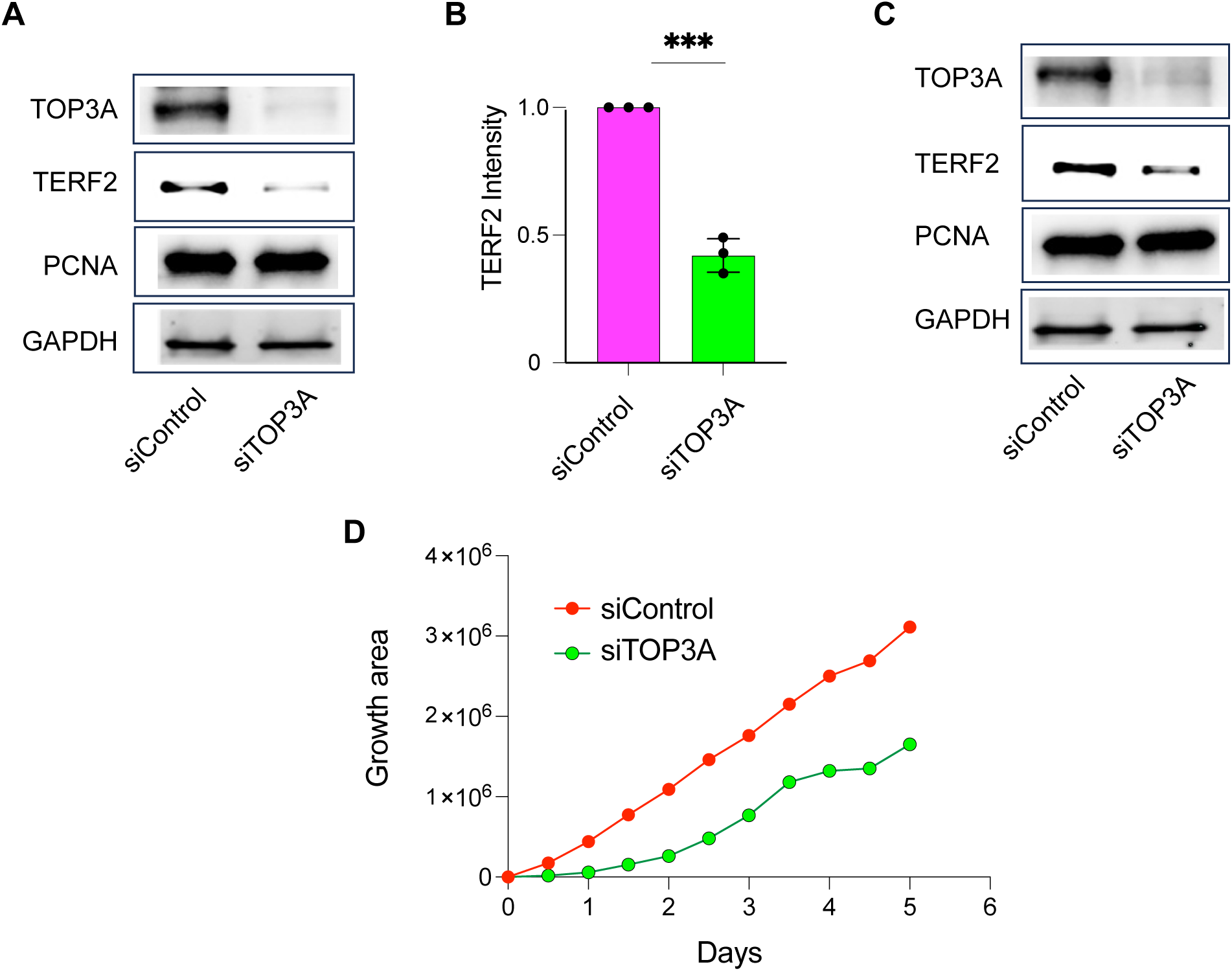
**A.** TERF2 immunoblot after TOP3A knockdown in SAOS2 (ALT) cells. **B.** Quantification of TERF2 intensity using image J software and normalization with GAPDH in SAOS2 cells. The experiment was performed in triplicates and average TERF2 intensity calculated. **C.** TERF2 immunoblot after TOP3A knockdown in HU09 (ALT) cells. **D.** Growth of SAOS2 cells was monitored after knockdown of TOP3A using Bio-spa. The experiment was done in duplicates.

**Supplementary Table S1:** List of important chemicals and Reagents.

| **Antibodies** | **Company** | **Catalog** |
| --- | --- | --- |
| Anti terf2 | Sigma millipore | 05-521 |
| Anti terf2 | Novus Biologicals | NB 110-57130 |
| Anti TOP3A | Proteintech | 14525-1-AP |
| Anti PML | Santacruz Biotechnology | SC-966 |
| Anti BLM | Bethyl laboratories | A300-110A |
| Anti TOP3A | *Smaali N T, EMBO J, 2008* |  |
| Anti PCNA | Santacruz Biotechnology | SC-56 |
| Anti GAPDH | Cell signalling | 14c10 |
| **PNA probe** |  |  |
| TelG-Cy3 | PNA bio | F1006 |
| TelC-Alexa488 | PNA bio | F1004 |
| **Plasmid** |  |  |
| GFP-TERF2 | Origene | RG223601 |
| HALO-TOP3A | This study |  |
| FLAG-TOP3A | *Saha L, Nature communication, 2023* |  |
| FLAG-TOP3A R36W | *Saha L, Nature communication, 2023* |  |
| **Important Chemicals** |  |  |
| Cycloheximide | Sigma Aldrich | 239763-M |
| Tris HCL pH7.5 | KD medical | 50-103-1383 |
| BSA | Sigma Aldrich | A2058 |
| Goat Serum | Sigma Aldrich | G9023 |
| DMEM | Thermofisher scientific | 11965-092 |
| RPMI 1640 | Thermofisher scientific | 11875-093 |
| RNAimax transfection kit | Invitrogen | 56532 |
| Trypsin | Thermofisher scientific | 25200056 |
| Blocking reagent | Roche | 11096176001 |
| 20X SSC buffer | Invitrogen | 15557-036 |

